# Shapes of Plasma Membrane Vesicles and PC-Cholesterol Vesicles Reveal Their Effective Spontaneous Curvature

**DOI:** 10.1101/2024.11.04.622000

**Authors:** Harshmeet Kaur, Tanmay Pandey, Tripta Bhatia

## Abstract

Giant membrane vesicles (GUVs) and giant plasma membrane vesicles (GPMVs) are useful models for studying cellular membrane properties. Our research analyzed the reduced volume of vesicles made from phospholipid and 10% cholesterol to investigate transbilayer sugar asymmetries. We found that GPMVs have an average reduced volume of (0.88 *±* 0.06) with buffer asymmetry of 323 mM, lower than the (0.92 *±* 0.08) observed for DOPC: cholesterol vesicles with sucrose/glucose asymmetry of 390 mM. GUVs with different sugars inside and outside were more deflated, demonstrating a greater volume reduction than those with the same sugar inside and out. We applied the area-difference elasticity (ADE) model to map GPMVs and used the spontaneous curvature (SC) model to analyze DOPC: cholesterol GUVs, extracting spontaneous curvature based on their reduced volume.

**TOC Graphic:** 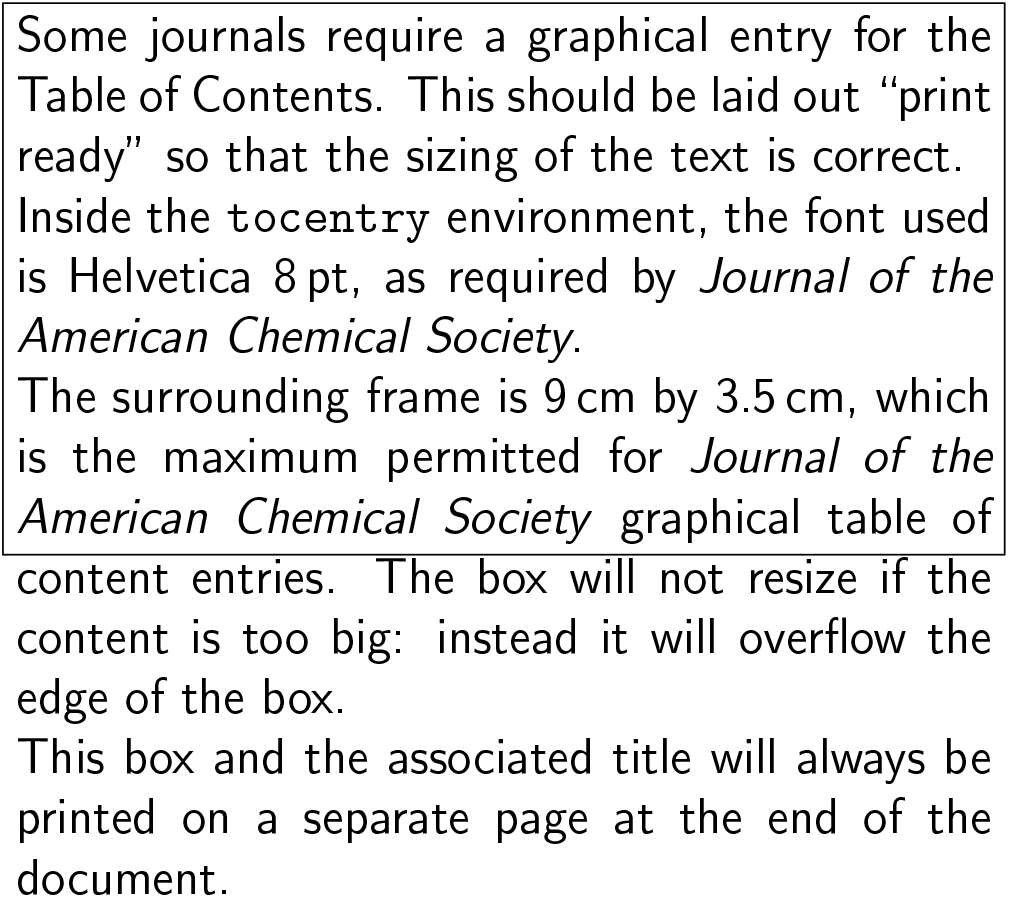

## Introduction

The plasma membrane separates the cell’s internal components from the outside environment, emphasizing the key role of lipid-protein membranes in the emergence of life. Primitive membranes likely consist of fatty acids and cholesterol, and their properties are influenced by environmental factors that shape the evolution of biological membranes. Giant plasma membrane vesicles (GPMVs) are model systems to study cell membranes. Figure 1 shows human embryonic kidney (HEK) 293 cells with GPMVs, which typically range from 5 to 20 micrometers (µm) in diameter.^1–3^ GPMVs are extracted directly from the plasma membrane of cells using the chemical induction method.^1^ Cells contain a cytoskeleton and complex lipid mixtures that differ between the cytoplasmic and extracellular leaflets of the plasma membrane. When giant plasma membrane vesicles (GPMVs) are extracted from HEK293 cells, the surface area of the two monolayers within the GPMVs differ in composition. GPMVs lack a cytoskeletal network, which normally provides shear elasticity by anchoring specific protein complexes to the membrane from the cytoplasmic side. In GPMVs, both solution asymmetry and compositional asymmetry are present, and these factors work together to influence the shape and physicochemical properties of the vesicles.

**Figure 1.**
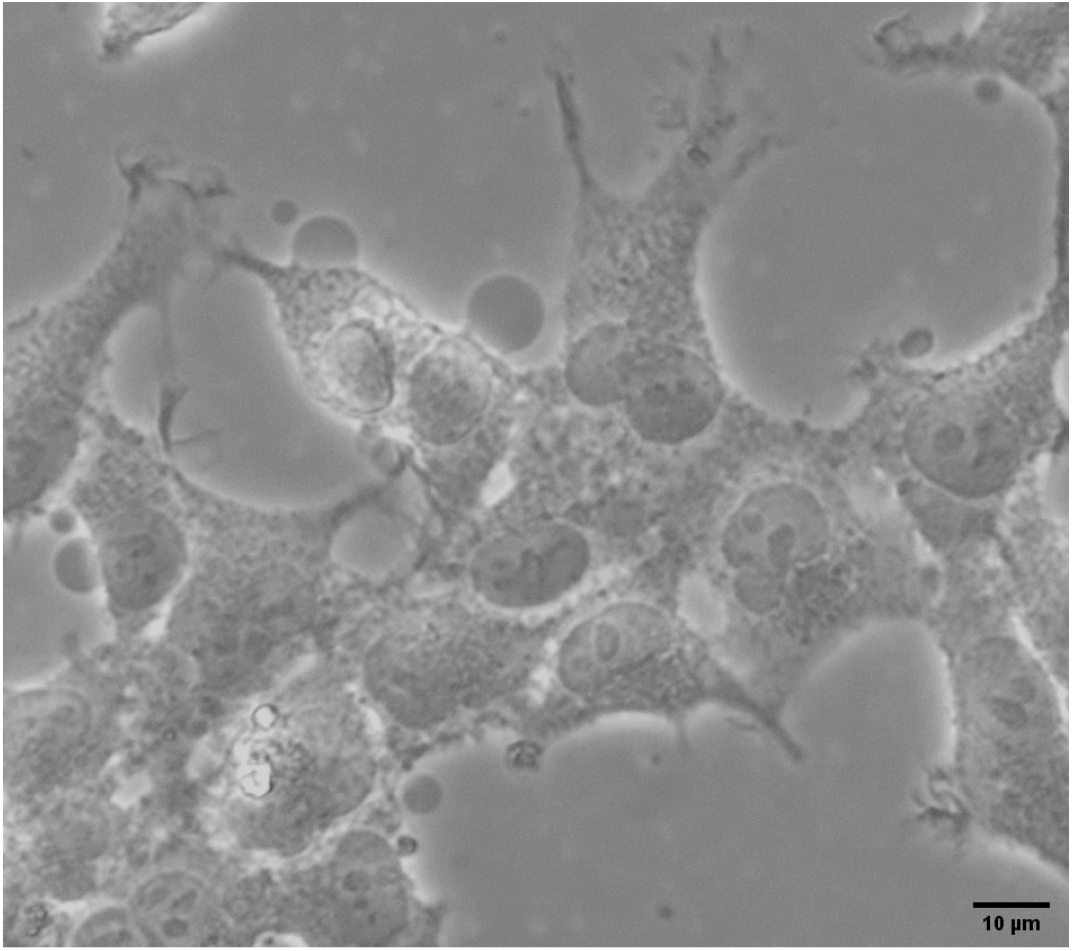
The human embryonic kidney 293 cells were visualized with Giant plasma membrane vesicles using phase contrast microscopy with a 40X objective.

This paper explores the effective spontaneous curvature and reduced volume, which are key to understanding the lipid and protein compositions in giant plasma membrane vesicles (GPMVs) and giant unilamellar vesicles (GUVs) made of DOPC and 10% cholesterol.

Figures 3a and 3b illustrate the effective spontaneous curvature and the spontaneous curvature, respectively, as functions of reduced volume in the stationary shape phase diagram discussed in the works of Döbereiner et al.^4^ and Seifert et al.^5^ respectively (reproduced with permission from the Journal). Using a precisely measured shape and a known reduced volume, we apply these parameters to a theoretically predicted catalog of stationary energy shapes in the phase diagram shown in Fig. 3 to predict membrane asymmetry.^4,5^ We used the area-difference elasticity (ADE) model, developed by Berndl, Kas, Lipowsky, Sackmann, and Seifert, to map the shapes of Giant Plasma Membrane Vesicles (GPMVs) onto the model’s phase diagram, allowing us to determine the effective spontaneous curvature 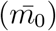 based on reduced volume.^5–11^

In contrast, Miao, Fourcade, Rao, Wortis, Zia, and Seifert focused on a preferred radius of curvature in their spontaneous curvature (SC) model, which overlooks the membrane’s bilayer nature.^5–11^ We applied the SC model to analyze Giant Unilamellar Vesicles (GUVs) made of DOPC: cholesterol and extract the spontaneous curvature (*c*_0_) from the given reduced volume. GUVs are spherical vesicles that consist of a single lipid bilayer and are formed using the electroformation, ^12^ gel-assisted method,^13^ or extrusion methods.

## Experimental

### Materials

We purchased powdered lipids from Avanti Polar Lipids. We dissolve the lipid powder in chloroform and keep it at −20°*C* for storage. We have used 1,2-Dioleoyl-sn-glycero-3-phosphocholine lipids (DOPC) (catalog no. 850375P) and cholesterol lipids to make vesicles. For making a buffer solution, Sucrose (purchased from Emplura) and glucose purchased from Sigma-Aldrich (CAS no. 50-99-7) were used. Milli-Q water is used as a solvent to make a buffer solution. For preparing GPMVs, we used NaCl (from Sigma-Aldrich, CAS no. 7647-14-5), Calcium chloride (from Avra, Catalog No. ASC2461), HEPES (from Sigma-Aldrich, H3375), KOH (from SRL, CAS no. 1310-58-3), dithiothreitol (DTT) (CAS no. 3483-12-3, trypsin and formaldehyde (HCHO)(CAS no. 50-00-0) are purchased from Himedia, HEK293 cell line (epithelial cells from the kidney of a human embryo that have adherent properties) purchased from ATCC (CAT no. CRL-1573).

### Preparation of GUVs

The electroformation method is used for the production of giant unilamellar vesicles. In this method, lipids are coated on two glass plates with indium tin oxide (ITO), and then chloroform is evaporated in a vacuum. Putting the Teflon spacer between two ITO slides coated with lipids, the lipids face each other, creating a chamber. The buffer solution was filled into the chamber with a syringe and connected to the function generator via copper tape. Two clamps were used to keep the ITO glass plates together to prevent any solution from escaping the cabinet. An AC sinusoidal field was applied to the electroswelling process. In the electroswelling process, peak voltages (200 − 2000) mV and frequencies (4 − 10) Hz were used in steps. DOPC: cholesterol (10%) GUVs were prepared in 300 *m*M sucrose inside the GUV and were transferred in the same or different concentrations of glucose outside the GUV, thereby generating a sugar solution asymmetry across the membrane, as listed below and in Table 1.

**Table 1:** The equilibrium concentration of GUV samples is determined by several parameters. The first column lists sample labels for different experiments. The second column shows the solution concentration (*C*_*i*_) used for GUV/GPMVs preparation, while the third column presents the solution concentration (*C*_*o*_) for imaging. The fourth column indicates the differences in inside-out solution concentration (Δ*C*). *C*_*eq*_ represents the equilibrium solution concentration inside the GUVs after achieving osmotic balance. The last column identifies the interior (In) and exterior (Out) solutions.

| **DOPC: cholesterol (10%) GUVs** |  |  |  |  |  |
| --- | --- | --- | --- | --- | --- |
| | $C_i$ | $C_o$ | $\Delta C$ | $C_{eq}$ | In/Out |
| Sample no. | (mM) | (mM) | (mM) | (mM) |  |
| I | 300 | 300 | 0 | 300 | suc/glu |
| II | 300 | 350 | 50 | 330 | suc/glu |
| III | 300 | 400 | 100 | 360 | suc/glu |
| IV | 300 | 450 | 150 | 390 | suc/glu |
| **GPMVs derived from cell** |  |  |  |  |  |
| V | 280-300 | 355.5 | $\geq 55.5$ | 323 | buffer1/buffer2 |
| **DOPC: cholesterol (10%) GUVs** |  |  |  |  |  |
| VI | 300 | 300 | 0 | 300 | suc/suc |

- Sample I: 300 mM sucrose inside and 300 mM glucose outside.
- Sample II: 300 mM sucrose inside and 350 mM glucose outside.
- Sample III: 300 mM sucrose inside and 400 mM glucose outside.
- Sample IV: 300 mM sucrose inside and 450 mM glucose outside.
- Sample VI: 300 mM sucrose inside and outside.

### Preparation of GPMVs

GPMVs were isolated from the HEK-293 cell line. There are a large number of methods for the extraction of GPMV from adherent cells. In this experiment, the chemical induction method was used.^1^ Vesiculation buffer, consisting of 150 *m*M NaCl, 2 *m*M CaCl_2_ and 20 *m*M HEPES, was prepared, and the pH was maintained at 7.4 using a 1 M KOH solution. Then, the active vesiculation buffer, consisting of 1.9 *m*M DTT and 27.6 *m*M formaldehyde, was prepared using vesiculation (prepared above) as a solvent on the same day of extraction of GPMVs. Now, to isolate GPMVs, a 25 cm^2^ flask of adherent cells (HEK293) was taken, which was trypsinized one day earlier. The medium was removed from above the cells, and the cells adhered to the surface were washed with 2 *ml* of vesiculation buffer. Finally, 1 *ml* of active vesiculation buffer was added to the flask and kept in the incubator at 37° *C*, gently shaking the flask every 10 minutes for 1 hour 30 minutes.

The osmolality of the cytoplasm of cells is about 280-300 mOsm/kg (buffer 1). To calculate the total osmolality of the complete culture media consisting of 89% Gibco RPMI 1640 culture media, 10% FBS, and 1% antibiotic solution, we need to consider the contributions of each component. Thus, the total osmolality of the culture media (buffer 1) with 10% FBS, 1% antibiotic solution, and 89% RPMI 1640 is approximately 283.65 mOsm/kg. For GPMVs extraction, cell culture media was removed and we added the active vesiculation buffer (buffer 2) containing 150 *m*M NaCl, 2 *m*M CaCl_2_ and 20 *m*M HEPES, 1.9 *m*M DTT and 27.6 *m*M formaldehyde (HCHO) giving a total osmolality as 355.5 mOsmol/kg. Inside GPMVs, the osmolality is approximately 280-300 mOsm/kg, while outside GPMVs, it is about 355.5 mOsm/kg.

### Confocal microscopy

A confocal microscope (Zeiss Axio Observer LSM 980 Airyscan-2) with a Plan-Apochromat 63x/1.40 oil immersion objective was used for three-dimensional imaging of giant unilamellar vesicles (GUVs) and GPMVs. To create a *z*-stack for shape analysis, the *z* range was defined by determining the top and bottom of the sample, with a *z*-step size of 0.5 *µm*. The pinhole size was set to 5 AU (approximately 293 *µm*), and the scan area was set to 2x. Nile red dye was used for fluorescence, with excitation at 559 *nm* and emission at 636 *nm*.

## Results and Discussion

### Theory of vesicle shape (revisited)

The Helfrich free-energy describes vesicle shapes in equilibrium, a general formulation given by^14^

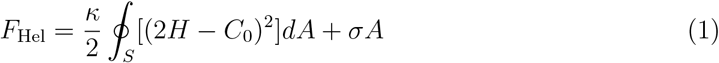

for a single vesicle with spherical topology. The material’s properties are defined by three parameters: the elastic modulus of bending (*κ*), the internal tension of the membrane (*σ*), and the spontaneous curvature (*C*_0_).^4,15–17^ Additionally, *H, A* represent the local mean curvature and the total surface area of the GUV, respectively. Let *N* ^out^ and *N* ^in^ be the number of lipid molecules in the outer and inner monolayers, respectively, with an area per lipid of *a*_0_ in the relaxed state. The geometric area difference Δ*A*_0_ between the two monolayers is defined by the difference in the number of lipid molecules.

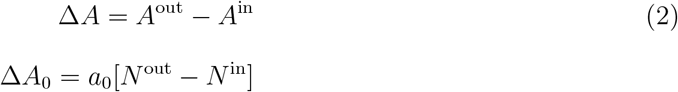

In Eq. 2, Δ*A* can vary due to several factors, such as changes in (i) *a*≠ *a*_0_, (ii) *N* ^out^, and (iii) *N* ^in^, resulting in (Δ*A* ≠ Δ*A*_0_). Cholesterol molecules can flip-flop from one leaflet to the second, changing *N* ^out^, and *N* ^in^ as well as effective *a*, in equilibrium, resulting in (Δ*A* = Δ*A*_0_). The volume of the vesicle *V* attains its maximal value when the vesicle has a spherical shape. The reduced volume (*v*)^11^ of the vesicle is defined as

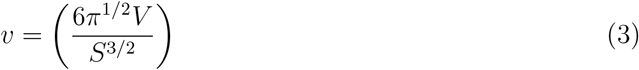

We captured z-stack sequences of vesicle outlines using confocal microscopy at a specific temperature. Our experiments focus on conditions where the *v* ≤ 1 allows for flaccid vesicles that can take on non-spherical shapes shown in Fig. 2. Table 2 presents the average value of the reduced volume 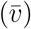 along with the error bar (Δ*v*) for the GUV and GPMV samples in osmotic equilibrium. The data is illustrated in the graph in Fig. 4. Solution asymmetries are found in samples I-V. In samples I-IV, sucrose is inside the giant unilamellar vesicles (GUVs), while glucose is in the external compartment. Sample V contains giant plasma membrane vesicles (GPMVs) from cells, with an internal buffer osmolality of 280-300 mOsm/kg and an external buffer osmolality of about 355.5 mOsm/kg. Sample VI has sucrose inside the DOPC: Cholesterol (10%) GUV and in the exterior compartment.

**Table 2:** The average value of the reduced volume 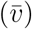 with the error bar (Δ*v*) in equilibrium for samples I-VI.

| Sample no. | $\bar{v} \pm \Delta v$ | Model | In/Out |
| --- | --- | --- | --- |
| I | $0.91 \pm 0.07$ | SC | Suc/Glu |
| II | $0.90 \pm 0.10$ | SC | Suc/Glu |
| III | $0.92 \pm 0.08$ | SC | Suc/Glu |
| IV | $0.87 \pm 0.12$ | SC | Suc/Glu |
| V | $0.88 \pm 0.06$ | ADE | buffer1/buffer2 |
| VI | $0.98 \pm 0.02$ | - | Suc/Suc |

**Figure 2.**
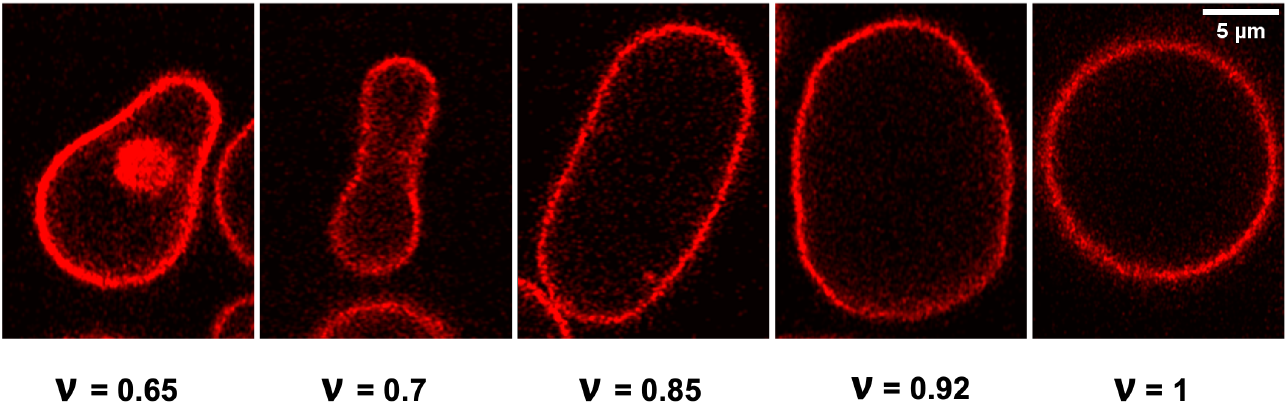
The experimental shapes of DOPC: cholesterol (10%) vesicles in osmotic equilibrium. The comparison illustrates the changes in the reduced volume (*v*) of DOPC: cholesterol (10%) vesicles when there is no concentration imbalance across the GUV membrane.

**Figure 3.**
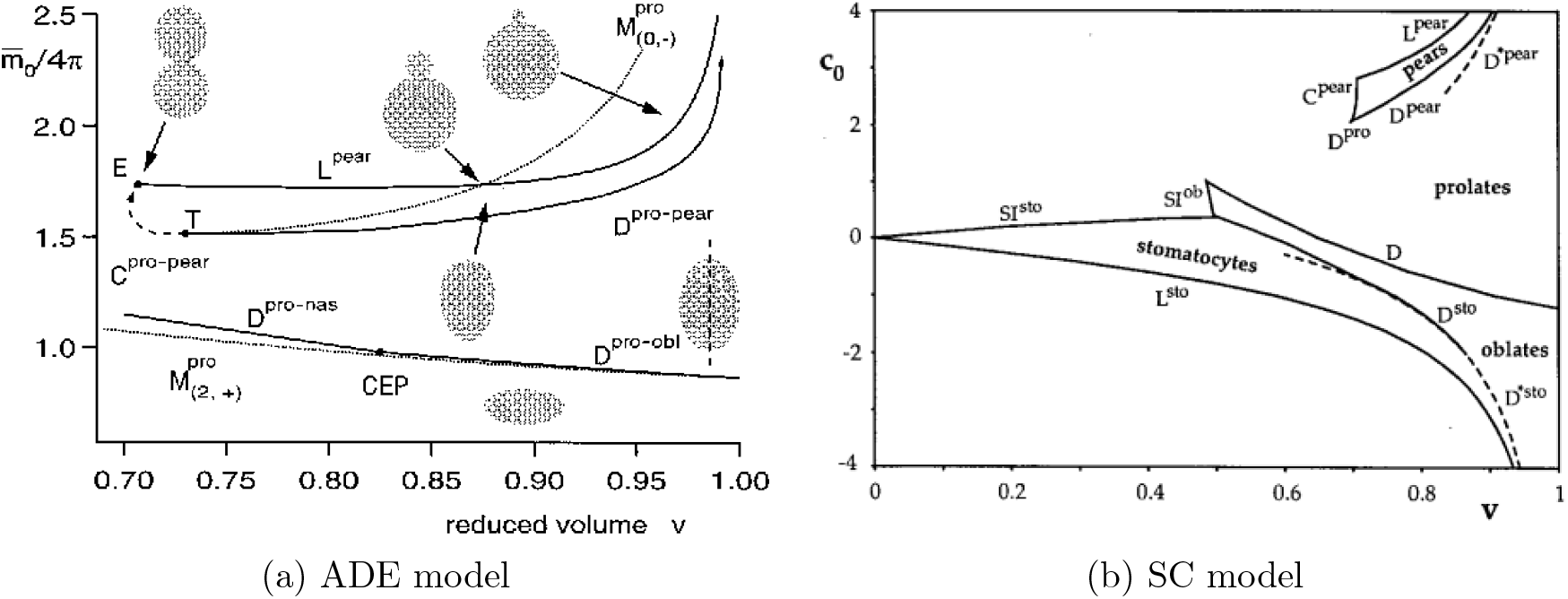
Phase diagrams reproduced from^4,5^ for the Lowest-energy shapes. (a) Leftside plot shows 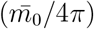 is plotted with the reduced volume for *α* = 1.4. Solid lines indicate first-order boundaries (D), second-order boundaries (C) by dashed lines, and spinodals (M) by dotted lines. (b) The right-side plot shows *c*_0_ is plotted with the reduced volume of the SC model.^5^

**Figure 4.**
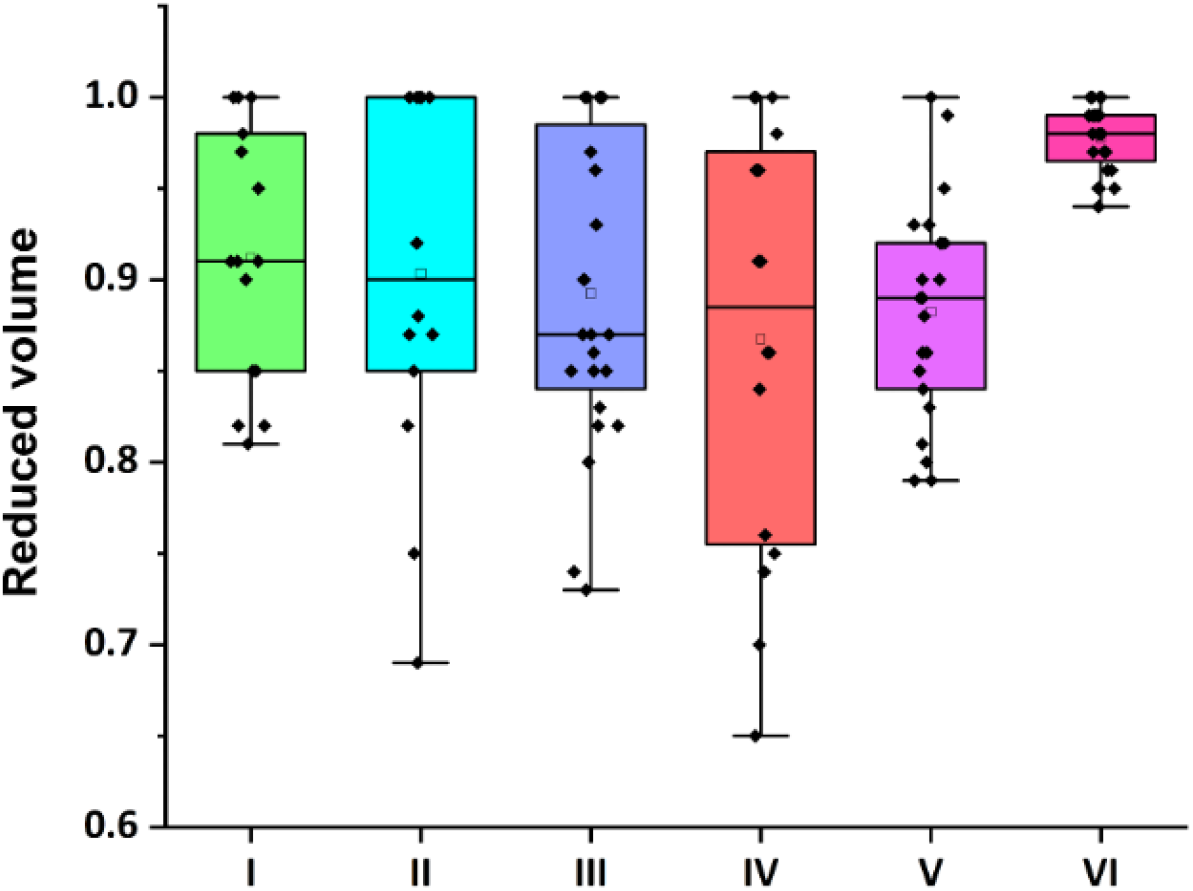
The comparison illustrates the changes in the reduced volume (*v*) of DOPC: cholesterol (10%) vesicles and GPMVs at equilibrium when there is no concentration imbalance across the GUV membrane. **I**: DOPC: Cholesterol (10%) GUVs (300 *m*M sucrose inside and 300 *m*M glucose outside), **II**: DOPC: Cholesterol (10%) GUVs (300 *m*M sucrose inside and 350 *m*M glucose outside) GUVs, **III**: DOPC: Cholesterol (10%) GUVs (300 *m*M sucrose inside and 400 *m*M glucose outside) GUVs, **IV**: DOPC: Cholesterol (10%) GUVs (300 *m*M sucrose inside and 450 *m*M glucose outside) GUVs, **V**: GPMVs, **VI**: DOPC: Cholesterol (10%) GUVs (300 *m*M sucrose inside and outside). The labeled I, II, III, IV, and V samples exhibit asymmetry in their solutions.

**Figure 5.**
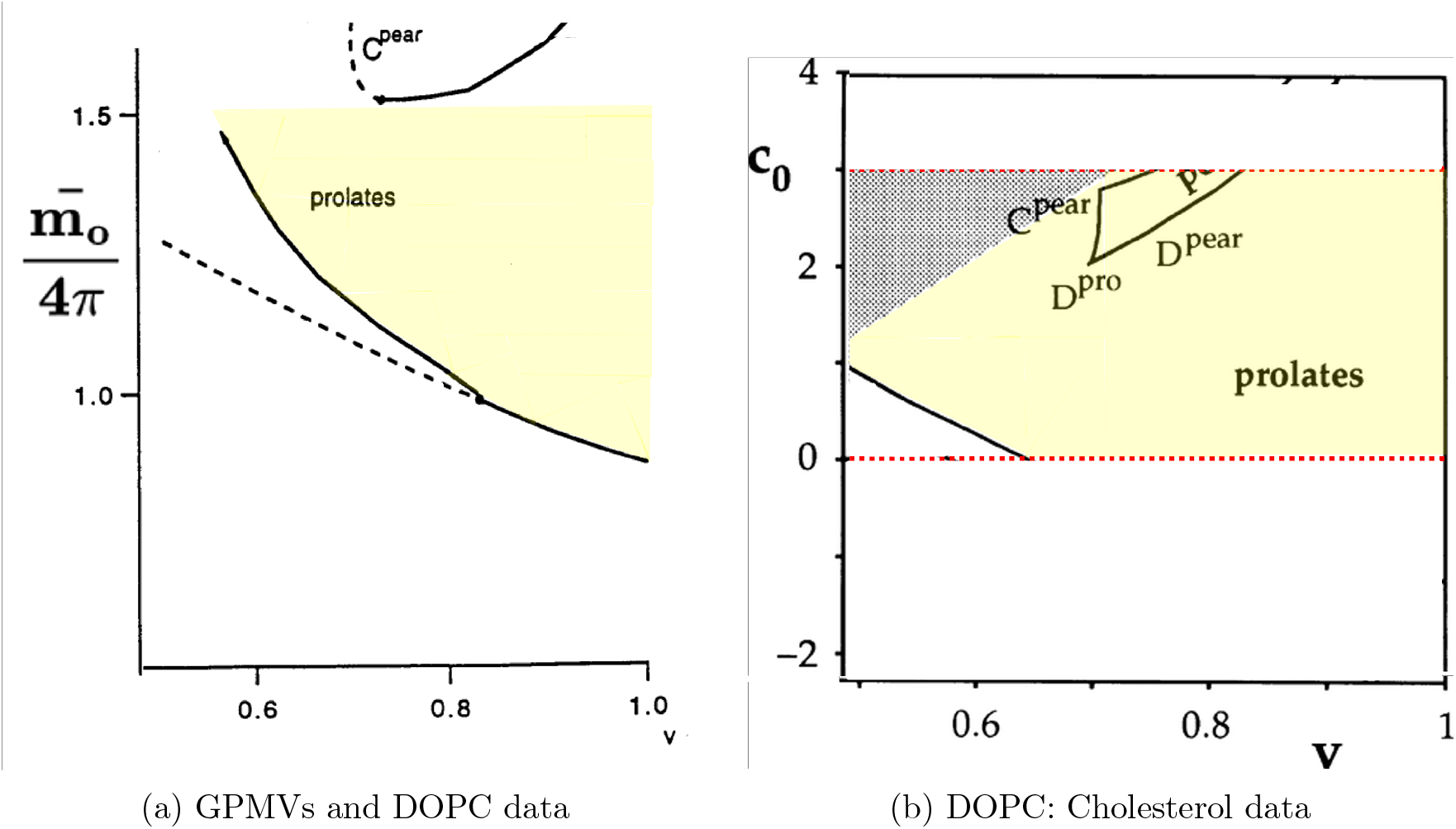
The yellow shaded region indicates the reduced volume for the observed shapes for(a) DOPC GUVs^22^ and GPMVs and for (b) DOPC: Cholesterol (10%) GUVs in the Phase diagrams reproduced from^4,5^ for the Lowest-energy shapes.

**The area-difference elasticity (ADE) model** estimates the elastic bending energy by integrating over the whole vesicle area for a given fixed enclosed volume (*V*), membrane area (*A*) and area-difference (Δ*A* − Δ*A*_0_) of the vesicle, using Eq. 4.

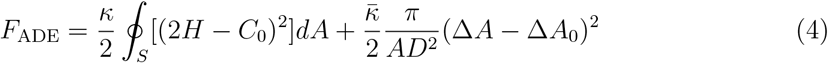

The ratio is 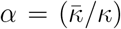 is the order unity for common phospholipids.^18^ The dimensionless area difference (reduced form) is defined as,

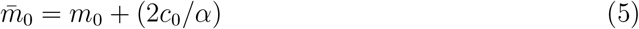

where for a fixed bilayer separation (*D*), the difference in geometrical area between the exterior and interior monolayers is (Δ*A*_0_), *R* is the radius of the GUV, *C*_0_ = 1*/R* is the spontaneous curvature, *c*_0_ = (*C*_0_*R*), is the dimensionless spontaneous curvature, *κ* is the bending modulus, and 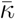 is the non-local bending modulus.,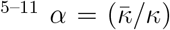 and *m*_0_ = (Δ*A*_0_*/*2*DR*). The spontaneous curvature (*C*_0_) reveals the preferred radius of curvature of the relaxed bilayer. Fig. 3a shows the value of the 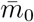 defined in Eq. 5 extracted from the stationary shape phase diagram discussed in the paper,^4^ reproduced with permission from the Journal. **The spontaneous curvature (SC) model** gives the elastic bending energy by integrating over the whole vesicle area, the first term in the Eq. 1 for given fixed enclosed volume (*V*) and membrane area (*A*) of the vesicle, assuming (Δ*A* = Δ*A*_0_). After the vesicle forms, the leaflet area can change in the samples discussed here because of the cholesterol flip-flop between the two monolayers of the bilayer. Thus, the difference in relaxed area (Δ*A* − Δ*A*_0_) between the two monolayers, which is based on the variation in the number of lipid molecules present, will be nearly zero, giving *m*_0_ = 0 in Eq. 5.^5–11^ Fig. 3b shows the value of the *c*_0_ defined in Eq. 5 extracted from the stationary shape phase diagram discussed in the paper Seifert et al.,^5^ reproduced with permission from the Journal.

### Experimental data for GPMVs and DOPC: Cholesterol vesicles

Using confocal images, we can calculate the surface area and volume of the vesicle, as well as the reduced volume. Fig. 4 shows reduced volume plots for samples I-VI (see Table 1). The second column in Table 2 shows the average value of the reduced volume 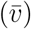 with the error bar (Δ*v*) of the GUV samples in osmotic equilibrium. Solution asymmetries are present in samples I -V. The giant unilamellar vesicle membrane can develop transbilayer asymmetry due to the uneven absorption of sugars. Glucose, which is smaller than sucrose and has additional chemical groups, can interact more effectively with the lipids in the outer leaflet, leading to densely packed adsorption layers.^16,19–21^ The plot with the reduced volume data is shown in Fig. 4 and the average reduced volume data is given in Table 2.

### Role of solution asymmetry in shape deflation

In response to any solution asymmetry, such as the differing composition of interior and exterior solutions in Table 1 for samples I-V, the quantitative analysis presented in our manuscript refers specifically to the equilibrium shapes achieved. From Table 1, we find that for DOPC: cholesterol (10%) GUVs without solution asymmetry across the membrane, deflation is less significant compared to the case with solution asymmetry. The average value of the reduced volume in the absence of the solution asymmetry (sample VI) is (0.98 *±* 0.02), shown in Table 1. The deflation of DOPC: cholesterol (10%) GUVs is relatively less, and the value of the average reduced volume data in Table 1 indicates that vesicles with different sugars inside and outside the GUV are more deflated than those with the same sugar inside and outside, which tend to be nearly spherical.

We also studied the GPMVs prepared by HEK293 cells ; the resulting reduced volume is lower than DOPC: Cholesterol vesicles GUVs without solution asymmetry. It is important to note that GPMVs have membrane asymmetry due to compositional heterogeneity and solution transbilayer asymmetry derived from the cellular plasma membranes. GPMVs derived from the cell’s plasma membrane have different buffer mediums inside and outside the membrane. In osmotic equilibrium, the GPMVs have an average value of reduced volume as 0.88 *±* 0.06, which is lower than 0.98 *±* 0.02 for DOPC: Cholesterol vesicles in the absence of sugar asymmetry, i.e., sucrose inside the GUV and in the exterior compartment of GUVs. However, GUVs with different sugars inside and outside the GUV (samples I-IV) were more deflated, and their volume reduction was larger than GUVs with the same sugar inside and outside the GUV (sample VI).

### Role of concentration imbalance in shape deflation

We prepare GUVs with DOPC and cholesterol (10%) in 300 *m*M of sucrose. The giant unilamellar vesicles were transferred in a glucose solution with osmolarity differences of 50 *m*M, 100 *m*M, and 150 *m*M outside the GUV. This was done to create a concentration gradient across the membrane, as shown in Table 1. The corresponding reduced volume was calculated for each condition after achieving osmotic equilibrium. The average value of the reduced volume (see in Table 2) varies between (0.87 *±* 0.12) and (0.92 *±* 0.08) for the deflation of DOPC: cholesterol (10%) GUVs (samples I-IV), with sugar asymmetry, i.e., sucrose inside the GUV and glucose in the exterior compartment of GUVs.

In response to any concentration imbalances, such as (*C*_i_ *< C*_o_) in Table 1 for samples II-V, water flows from the interior to the exterior compartment. This process results in a reduction of enclosed volume through osmotic deflation. However, the quantitative analysis presented in the main part of our manuscript refers specifically to the equilibrium shapes achieved once osmotic equilibrium has been reached. The value of *C*_eq_ in Table 1 represents the equilibrium concentration of solution inside and outside after osmotic balance across the membrane has been attained. For DOPC: Cholesterol GUV Fig 4 shows that with an increasing solution concentration imbalance across the membrane, the deflation of GUVs remains the same within the error bar.

## Conclusion

The neutral Giant Unilamellar Vesicle (GUV) samples were prepared using a mixture of DOPC and cholesterol. One sample (Sample VI) contained a sucrose solution both inside and outside the GUVs, while the other samples (Sample I-IV) had a sucrose solution inside and a glucose solution outside the GUVs. Additionally, Giant Plasma Membrane Vesicles (GPMVs), derived from the plasma membrane of cells, were prepared with different buffer media inside and outside the membrane (Sample V). Bilayer asymmetry in GUVs is achieved by exposing each leaflet to different sugar solutions. ^16,21^ Previous research also revealed multispherical shapes resulting from this sugar solution asymmetry. ^19,20^

Figs 2, 4 show the vesicles’ shape change and their reduced volume in the absence and presence of membrane asymmetry in osmotic equilibrium across the membrane. GUVs with different sugars inside and outside the GUV were more deflated, and their volume reduction was larger than GUVs with the same sugar inside and outside the GUV. We also investigated the giant plasma membrane vesicles (GPMVs) prepared from HEK293 cells. It’s important to highlight that GPMVs exhibit membrane asymmetry due to compositional heterogeneity and transbilayer asymmetry in the solution, reflecting characteristics of the cellular plasma membranes.

We have shown that DOPC GUV and DOPC: DOPG(10%) (sugar and glucose) are vesicles that deflate more with an increasing solution concentration imbalance.^22^ and this paper for DOPC: cholesterol GUVs. Our work provides a useful way to calculate the effective spontaneous curvature from the reduced volume for membrane compartments by mapping the experimental shapes into the phase diagram.

## Author Contribution

HK performed the DOPC/cholesterol GUVs and GPMVs experiments, and TP wrote the code for the reduced volume analysis and performed data analysis. TB designed the experiments and the code and wrote the manuscript.

## Acknowledgements

This research was conducted within the Indian Institute of Science Education and Research Mohali and supported by the Ramalingaswami grant of DBT (DBT/RLF/Re-entry/06/2020) and IC-12025(22)/1/2023-ICD-DBT.

## Supporting Information Available

The data for samples I through VI of each GUV and GPMV has been shared.

